# Effects of combined prenatal exposure to air pollution and maternal stress on social behavior and oxytocin and vasopressin systems in male and female mice

**DOI:** 10.1101/2025.09.04.672354

**Authors:** Maura C. Stoehr, Elise M. Martin, Joy T. Babalola, Jason Xue, Matthew J. Kern, Niki Y. Li, Madeline F. Winters, Sarvin Bhagwagar, Caroline J. Smith

## Abstract

Prenatal exposures to air pollution and maternal psychosocial stress are each associated with increased risk of neurodevelopmental disorders, including autism spectrum disorder (ASD) and epidemiological work suggests that concurrent exposure to these risk factors may be particularly harmful. This is important given that the same populations often bear the brunt of both toxicant and psychosocial stress burdens. Social impairments are a defining symptom in ASD. Previous work modeling combined prenatal exposure to diesel exhaust particles (DEP) and maternal stress (MS) in rodents has found male-biased social deficits in offspring, as well as changes to neuroimmune processes and the gut microbiome. However, the precise neural circuits on which these exposures converge to impact social behavior is unclear. Oxytocin (OXT) and vasopressin (AVP) are neuropeptides critical to the regulation of social behavior across species, signaling primarily at the oxytocin receptor (Oxtr) and vasopressin V1a receptor (V1aR) in the brain. Here, we hypothesized that OXT and/or AVP expression would be reduced in the brain following DEP/MS exposure. Following prenatal exposure to DEP/MS or the vehicle/control condition (VEH/CON), we measured maternal and offspring outcomes during the perinatal period, social and anxiety-like behavior during adolescence, OXT and AVP cell/fiber density and *Oxtr* and *Avpr1a* mRNA expression in early adulthood in several brain regions in both males and females. We observed a decrease in interaction time in DEP/MS males as compared to VEH/CON in the sociability assay and a decrease in social novelty preference in DEP/MS females as compared to VEH/CON. No effects of sex or treatment were observed on OXT or AVP cell number or fiber density in the hypothalamic regions assessed. However, numerous sex differences were observed in *Oxtr* and *Avpr1a* mRNA. Moreover, *Avpr1a* mRNA was significantly increased following DEP/MS exposure in the nucleus accumbens in both sexes and tended to increase in the dorsal hippocampus. Conversely, *Avpr1a* mRNA tended to decrease in the amygdala in both sexes following DEP/MS exposure. Together, these findings suggest that DEP/MS exposure has a stronger impact on female social behavior than previously observed. Moreover, while DEP/MS exposure does not appear to impact OXT or AVP expression in the brain, V1aR expression is modulated by DEP/MS exposure in several brain regions.

**Highlights:**

- Prenatal DEP/MS reduces social novelty preference in females
- Prenatal DEP/MS does not alter OXT or AVP cell number in the PVN
- Prenatal DEP/MS does not alter OXT or AVP fiber density in the LH, AH, or MPOA
- Prenatal DEP/MS increases *Avpr1a* mRNA in the NAc
- Prenatal DEP/MS tends to decrease *Avrp1a* mRNA in the AMY and increase in dHipp

## Introduction

Air pollution is a massive public health burden with over 7.3 billion people exposed to unsafe levels of air pollution globally (Fuller et al., 2022; Southerland et al., 2022; Rentschler & Leonova, 2023). Furthermore, air pollution disproportionately impacts low-income communities that face higher levels of psychosocial stress and lower resource availability as compared to more affluent communities (Jbaily et al., 2022; Earnshaw et al., 2013). Both air pollution and maternal psychosocial stress during pregnancy are risk factors for neurodevelopmental disorders, such as autism spectrum disorder (ASD; Lyall et al., 2014; Dutheil et al., 2021; Volk et al., 2011; Volk et al., 2013; Roberts et al., 2014; Kinney et al., 2008a; Kinney et al., 2008b). Cumulative developmental exposures to air pollution and psychosocial stressors may have compounding effects (Margolis et al., 2025; Jáni et al., 2024). For example, McGuinn et al. (2019) found that PM_2.5_ exposure during the first year of life was associated with increased ASD risk and that association was moderated by neighborhood deprivation score (McGuinn et al., 2019). Similarly, maternal perceived stress is associated with decreased hippocampal volume and visuospatial reasoning in 5–7-year-old children, and this relationship is moderated by prenatal polycyclic aromatic hydrocarbon (a component of air pollution) exposure (Margolis et al., 2022; McGuinn et al., 2019). Taken together, the existing literature suggests that prenatal exposure to air pollution and maternal psychosocial stress may interact to alter neurodevelopmental trajectories in ways that may increase risk for neurodevelopmental and/or psychiatric disorders (Herting et al., 2024; Margolis et al., 2022; Johnson et al., 2021; Rivas et al., 2019).

Animal models lend further insight into the neurobiological mechanisms underlying disrupted neurodevelopment following prenatal toxicant and/or maternal stress exposures. Several studies using rodent models of air pollution and/or maternal stress during early postnatal development have observed changes in social behavior in offspring (Sobolewski et al., 2018; Zhou et al., 2021; Fonken et al., 2011; Li et al., 2018; Nephew et al., 2020; Weitekamp & Hofmann, 2021; Gur et al., 2019; Chen et al., 2024; Weinstock, 2016; Ehrlich & Rainnie, 2015; Lee et al., 2007; Church et al., 2018; Chang et al., 2018). In humans, disrupted social behavior is a defining symptom of ASD (American Psychiatric Association, 2022). Exposure to ultrafine particulate matter (UFP) during the first week of postnatal life decreased time spent exploring a novel social stimulus by adult male but not female mice (Sobolewski et al., 2018). Male rats exposed to PM_2.5_ from postnatal day (P)8-22 display decreased ultrasonic vocalizations, social interaction and social discrimination, and increased anxiety-like behavior as compared to controls (Li et al., 2018). Similarly, developmental exposure to particulate matter until the end of the lactation period decreased social play and allogrooming in adolescent male rats (Nephew et al., 2020). Importantly, these studies all involved air pollution exposure directly to individuals during the postnatal period. This postnatal exposure is fundamentally different from maternal gestational exposure which indirectly impacts offspring in utero via consequences of maternal immune activation (MIA; Block et al., 2022), rather than the immune sequelae that would result from direct air pollution exposure to offspring. In a line of work using a mouse model of combined exposure to diesel exhaust particles (DEP) and maternal stress (MS) during pregnancy (Bolton et al., 2013; Block et al., 2022; Smith et al., 2023), offspring are exposed to diesel exhaust particles throughout pregnancy and a mild form of maternal resource limitation stress during the prenatal, but not the postnatal period. In these studies, P7-P9 male and female offspring display increased ultrasonic vocalization (USV) frequency but reduced USV complexity following DEP/MS as compared to control mice (Block et al., 2022). During adolescence, DEP/MS males, but not females, exhibit deficits in sociability and social novelty preference behavior as compared to controls (Block et al., 2022; Smith et al., 2023). In adulthood, DEP/MS offspring also display impaired contextual fear recall, appetitive social behavior, courtship behavior, and functional activation of social behavior circuits in males only (Bolton et al., 2013; Block et al., 2022). Importantly, deficits in sociability and social novelty preference are not observed following either prenatal air pollution or maternal stress exposure alone, suggesting that the addition of a mild stress to mothers unmasks vulnerability to environmental toxicants (Smith et al., 2023), in line with findings from the human literature. However, which social behavior circuits are disrupted following DEP/MS behavior remains unknown.

The social decision-making network (SDMN) is a complex set of interconnected brain regions crucial to the generation of social behaviors that includes brain regions such as the nucleus accumbens (NAc), lateral septum (LS), hippocampus, and amygdala (AMY; O’Connell & Hofmann, 2011; O’Connell & Hofmann, 2012). Oxytocin (OXT) and vasopressin (AVP) are critical neuropeptide modulators of social behavior across species. OXT signaling has been implicated in numerous behaviors including maternal behavior, reproductive behavior, social investigation and recognition, social novelty-seeking, social avoidance, cooperative behaviors, and social play, often with sex-specific effects (for recent reviews see Procyshyn et al., 2024; Menon & Neumann, 2023; Rigney et al., 2022). Meanwhile, AVP signaling regulates aggression, social recognition, cooperative behaviors, partner preference, social communication, social play, social investigation, and others (for recent reviews see Rigney et al., 2022; Rigney et al., 2023; Kompier et al., 2019). Both OXT and AVP are synthesized in hypothalamic nuclei, most largely the paraventricular nucleus of the hypothalamus (PVN) and the supraoptic nucleus of the hypothalamus (SON; Veenema & Neumann, 2008; Buijs et al., 1983; Landgraf & Neumann, 2004) and in the extended amygdala (De Vries & Panzica, 2006; Rood & De Vries, 2011; Young & Gainer, 2003; Lee et al., 2009; Liao et al., 2020). Importantly, there is abundant expression of OXT and AVP fibers, as well as their respective receptors: the oxytocin receptor (Oxtr) and the vasopressin 1a receptor (V1aR) throughout the regions of the SDMN (Veenema & Neumann, 2008; Tan et al., 2019; Caldwell, 2017; Bredewold et al., 2018; Neumann & Landgraf, 2012; Albers, 2012; Caldwell et al., 2008; Smith et al., 2017; DiBenedictis et al., 2017; Wang et al., 1996; Albers, 2015; De Vries & Panzica, 2006; Rood & De Vries, 2011; Newmaster et al., 2020; Dumais et al., 2013; Insel et al., 1991; Insel et al., 1993; Olazaábal & Young, 2006; Lee et al., 2008; Inoue et al., 2022).

There is a wealth of literature examining the effects of perinatal environmental toxicant exposures on OXT and AVP systems, particularly plastics, flame retardants, and pesticides (for reviews see Martin et al., 2025; Patisaul, 2017). However, there is very little work on the effects of perinatal air pollutants on OXT, AVP, and their respective receptors in rodent models. DEP/MS induces MIA (Block et al., 2022) and several studies using MIA models have found changes in OXT, Oxtr, and AVP (Taylor et al, 2012; Breach et al., 2024; Goh et al., 2020; Ronovsky et al., 2017). Similarly, studies of maternal stress during pregnancy have found decreased *Oxtr* mRNA expression in the prefrontal cortex of male offspring (Gur et al., 2019). However, it is unknown how combined exposures to air pollution and maternal stress impact OXT and AVP cell and receptor expression in the brain.

For the present study, we aimed to assess the impact of prenatal DEP/MS on social behavior, OXT and AVP cells and their projections in the hypothalamus, and *Oxtr* and *Avpr1a* mRNA expression within regions of the SDMN as compared to VEH/CON in both males and female offspring. Specifically, we measured OXT-immunoreactive (-ir) and AVP-ir cell number and fiber density within the PVN, lateral hypothalamus (LH), anterior hypothalamus (AH), and medial preoptic area (MPOA). We assessed *Oxtr* and *Avpr1a* mRNA expression in the NAc, LS, AMY, dorsal hippocampus (dHipp), and ventral hippocampus (vHipp) following DEP/MS in male and female offspring. Based on the previous work in this model that reported male-specific decreases in social behavior (Block et al., 2022; Smith et al., 2023), we hypothesized that DEP/MS would reduce the expression of OXT and/or AVP system parameters in many of these regions in male offspring, corresponding to social deficits.

## Methods

### Animals

Wild-type (WT) C57Bl/6J mice were purchased from Jackson Laboratories and bred in the laboratory for two generations before use in experiments. Animals were group-housed with same-sex littermates under standard laboratory conditions (12-hour light/dark cycle [6am-6pm], 68° F, 50% humidity, *ad libitum* access to food and water) prior to experiments. To account for litter effects, offspring were obtained from multiple litters (1-4 pups per sex per litter from eight litters total) in all treatment groups. Experiments were conducted following the NIH *Guide to the Care and Use of Laboratory Animals* and approved by the Institutional Animal Care and Use Committee (IACUC) at Boston College.

### DEP/MS Exposures

#### DEP Instillations

DEP was obtained from Dr. Staci Bilbo at Duke University. These particles originate from Dr. Ian Gilmour at the Environmental Protection Agency and are consistent in composition to those used in previous studies (Bolton et al., 2013; Bolton et al., 2017; Block et al., 2022; Smith et al., 2023). Instillation procedures were conducted in accordance with previously published work using this model (**Figure 1A**; Bolton et al., 2013; Block et al., 2022; Smith et al., 2023). Nulliparous C57Bl/6J females were time mated and the presence of a vaginal plug was used as an indication of pregnancy and set as embryonic day (E)0. Females were then pair-housed based on embryonic day until E13, at which point they were singly housed in cages containing a thin layer of AlphaDri bedding (AlphaDri; Shepherd Specialty Papers). Beginning on E2, dams received instillations every 3 days throughout pregnancy, totaling 6 instillations. For each instillation, females were weighed and anesthetized with 4% isoflurane. Dams were then suspended by their incisors from a plastic wire and received either 50μg/μL of DEP in vehicle (VEH; 0.05% Tween20 in phosphate buffered saline [PBS]) or VEH alone via oropharyngeal instillation over the course of 60s. Females were monitored in the home cage until they regained consciousness and resumed typical behavior (<5 min.).

**Figure 1.**
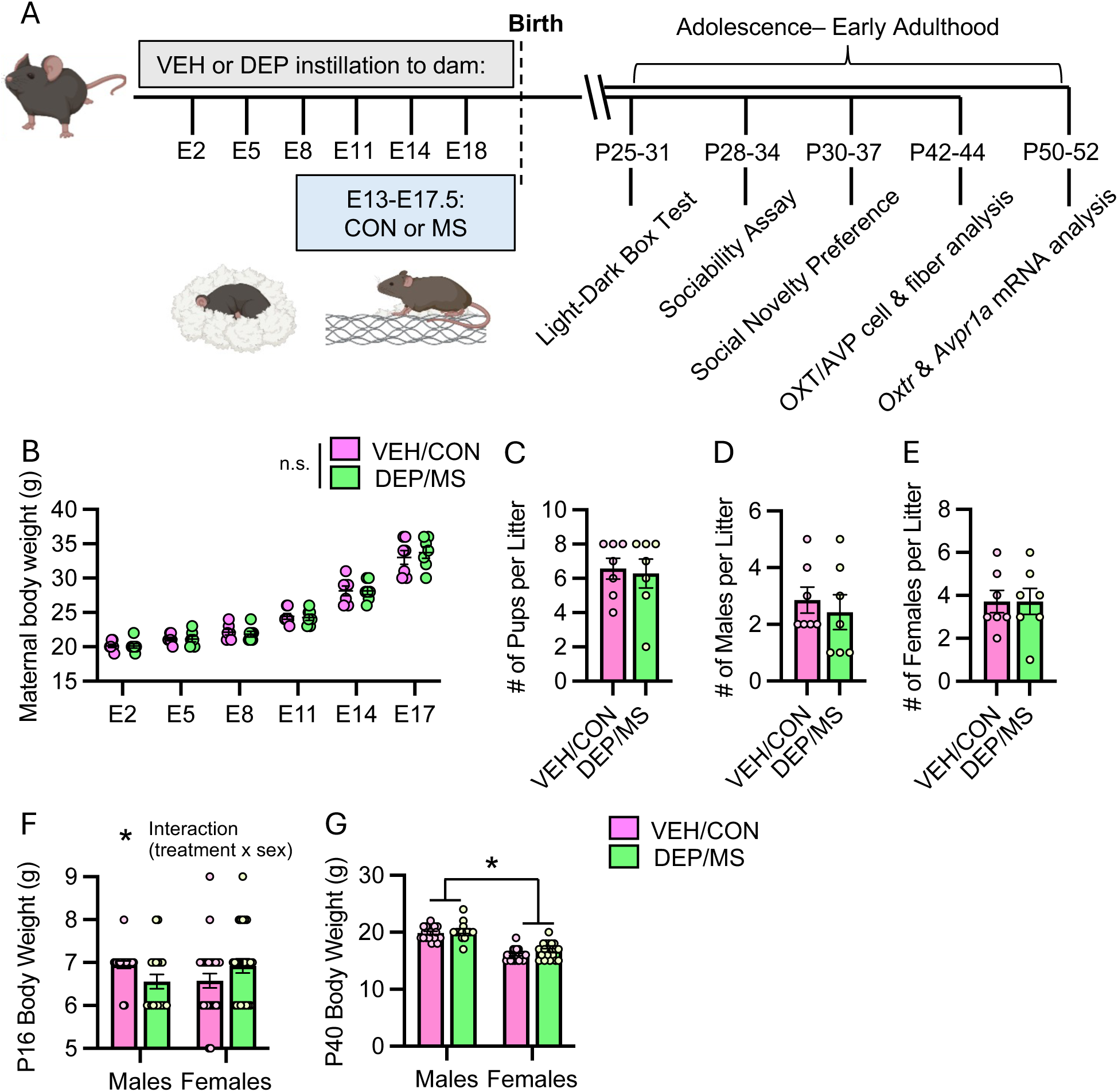
Perinatal following prenatal DEP/MS exposure. **A**) Schematic of experimental timeline. B) No effect of treatment was observed in maternal weight gain during gestation. **C**-**E**) No effects of DEP/MS treatment on overall litter size or number of male or female pups per litter as compared to VEH/CON. **F**) At P16, there is an interaction between sex and treatment on pup body weight, with DEP/MS males weighing less than VEH/CON males and DEP/MS females weighing more than VEH/CON females. **G**) At P40, there is a main effect of sex, but not treatment, with males weighing more than females. **B, F, G**: 2-way ANOVA (treatment x embryonic day or sex), **C-E**, unpaired t-tests. VEH = vehicle. CON = control bedding condition. DEP = diesel exhaust particles. MS = maternal stress condition. DEP = diesel exhaust particles. P = postnatal day. E=embryonic day. Data = mean ± SEM, *p<0.05.

#### Maternal Stress (MS)

To induce a mild maternal stressor (again in line with previous work using this model; Bolton et al., 2013; Block et al., 2022; Smith et al., 2023), during the last week of pregnancy (beginning on E13) females in the DEP condition were singly-housed in a cage containing a thin layer of AlphaDri bedding covered by an elevated aluminum mesh platform (0.4 cm x 0.9 cm mesh; McNichols Co., Tampa, FL) and given 2/3 (∼1.9g) of a cotton nestlet (MS condition). At the same timepoint, females receiving VEH instillations were placed into a cage with AlphaDri bedding and an entire nestlet (CON condition). On E17.5 (evening before delivery), all dams, regardless of treatment, were transferred to a clean cage of AlphaDri bedding with a full nestlet. Litters were left undisturbed during the first week of life. Litter size and sex ratios were recorded between postnatal day (P)7-10, and pup body weights were measured on P16 and P40. Offspring were weaned into cages with same-sex littermates at P24.

### Behavioral Testing

During the adolescent period (P25-P37), offspring underwent a series of behavioral assays each conducted during the second half of the light phase (1-5pm). Behavior was assessed sequentially on different days in the light-dark box test, the three-chambered sociability assay, and the three-chambered social novelty preference test. For all assays, males and females were tested using separate apparatuses on different days and each apparatus was disinfected between each test. Animals were handled and habituated to the testing room and the testing apparatus the day before testing. On testing days, animals were moved to the testing room 1hr prior to the beginning of the test to acclimate to the room. All videos were scored using Solomon Coder (Solomon.andraspeter.com) by blinded observers.

#### Light-Dark Box Test

The light-dark box is a square apparatus (40cm x 40cm) divided in half by a barrier with a small opening for animals to pass through (Figure 3A). One half of the box is made of clear plexiglass (the ‘light’ side) and the other half is opaque black and has a lid over the top (the ‘dark’ side). Mice were placed into the dark side of the box and allowed to freely roam between both sides of the box for 5 min. Time spent on the light side (s), dark side (s), and head-poking behavior (defined as the mouse’s head being on one side of the chamber while the rest of its body was on the other side; s), were quantified. Fecal boli deposition (N) was also quantified.

#### Sociability Assay

The three-chambered social preference test was performed according to Smith et al., (2023) and consists of a three-chambered arena (60cm x 40cm) with openings allowing passage between the chambers. Stimuli were confined within smaller containers (Plexiglass rod sides; diameter 10cm) in each of the side chambers. Subjects were placed into the middle chamber and freely allowed to investigate each stimulus (either a novel age-, sex-, and treatment-matched conspecific or a novel rubber duck) over the course of 5 min. Stimulus animals were habituated to the testing arena 1 day prior to testing. Behaviors quantified included: social, middle, and object chamber times (s), social and object stimulus interaction times (s; defined as time spent investigating/nose-poking into the stimulus container and climbing on it); social and object investigation time (s; nose-poking specifically), and social and object stimulus climbing (s; climbing specifically on the stimulus container rather than the chamber more generally). Interaction time included both nose-poking and stimulus climbing which were also scored separately. Middle chamber entries were recorded as a measure of locomotor activity. Animals were excluded from the test if they spent more than 25% of the test time climbing/sitting on top of the smaller cylinders containing the stimuli as this prevents them from engaging with the task (3 mice).

#### Social Novelty Preference Test

In the social novelty preference test, subject animals were placed in the middle of the three-chamber apparatus used for the sociability assay and allowed to freely investigate each stimulus (either a novel age-, sex-, and treatment-matched conspecific or a familiar age-, sex-, and treatment-matched cage mate) over the course of 5 min. Behaviors quantified included novel, middle, and familiar chamber times (s), novel and familiar interaction times (s; defined as time spent investigating/nose-poking into the stimulus container and climbing on it, novel and familiar investigation time (s; nose-poking specifically), and novel and familiar stimulus climbing (s; climbing specifically on the stimulus container rather than the chamber more generally). Interaction time included both nose-poking and stimulus climbing which were also scored separately.

### Immunohistochemistry (IHC)

#### Tissue Collection

For all IHC, animals were euthanized via CO_2_ inhalation between P42-P44 and brains were perfused with ice-cold saline. Brains were then removed and post-fixed for 48h in 4% paraformaldehyde, followed by 48h in 30% sucrose. Brains were then flash frozen in 2-methylbutane and stored at -20° C until sectioning at 30 μm on a cryostat (Leica Biosystems).

#### OXT and AVP staining in the hypothalamus

Free-floating sections containing hypothalamic nuclei (Allen Mouse Brain Atlas: https://atlas.brain-map.org; Bregma 0.02mm-1.06mm) were rinsed 5x in PBS, incubated in 10 mM sodium citrate for 30 min at 60° C for antigen retrieval, and rinsed 3x in PBS again. They were then incubated in a 1mg/1mL solution of sodium tetraborate in PBS at room temperature for 30 min, followed by 3x PBS rinses. Next, sections were blocked for 1hr in PBS with 10% normal goat serum, 0.3% Triton-X, and 1% H_2_O_2_. Sections were then incubated overnight with primary antibodies (guinea pig anti-OXT, 1:2000, Synaptic Systems; rabbit anti-AVP, 1:2000, Millipore Sigma) at RT on a shaker. After 3x PBS rinses, sections were then incubated in secondary antibodies (488 goat anti-rabbit IgG, 1:500, Invitrogen; 568 goat anti-guinea pig IgG, 1:500, Invitrogen) for 5 hr on a shaker, followed by 3x PBS rinses. Sections were mounted on gelatinized slides and cover slipped with Vectashield Antifade Mounting Medium with DAPI (Vector Laboratories).

#### Imaging

A Zeiss AxioImager Z2 microscope was used to take 10X magnification, tile-scanned images of the hypothalamus, including the PVN, LH, AH, and the MPOA. A 3x3 tile was used to capture the regions of interest. 7-step z-stacks were taken with a step-size of 1.53 μm measured from the center of the image to span 10 μm. Images were taken centering the third ventricle.

#### Quantification of OXT and AVP cell bodies in the PVN

Analysis of OXT-ir and AVP-ir cell bodies in the PVN was categorized based on OXT and AVP staining patterns. For OXT-ir quantification, sections were categorized into Level 1 (Bregma - 0.38mm to-0.56mm; the more anterior portion of the PVN with only sparse OXT-ir cell bodies) and Level 2 (Bregma -0.56mm to-0.88mm: dense OXT-ir cell bodies in heart-shaped formation). For AVP quantification, sections were also categorized into two Levels; Level 1 (Bregma - 0.28mm to -0.56mm: moderately dense AVP-ir cell bodies extending down along the third ventricle) and Level 2 (Bregma -0.56mm to -0.66mm: densest AVP-ir cell bodies in heart shaped formation. The expression patterns of OXT and AVP differ across the PVN (OXT expression is shifted slightly earlier in the PVN), thus, Level 1 is shifted slightly later for AVP. Z-stacks were converted into maximum intensity projections and cell bodies for OXT and AVP were hand counted in FIJI software (https://imagej.net/software/fiji) by blinded observers. For each stain (OXT vs. AVP) cell bodies were counted bilaterally in each coronal section and values for each level of the PVN for a given animal were averaged to provide a single measurement per animal.

#### Quantification of OXT and AVP fibers and/or cells in the LH, AH, and MPOA

Using the same images in which cell bodies were quantified in the PVN, OXT-ir and AVP-ir fibers were quantified in the LH (Bregma -0.48mm to -1.06mm), AH (Bregma -0.48mm to - 0.96mm), and MPOA (Bregma 0.02mm to -0.28mm) using FIJI. Regions of interest for each hypothalamic nucleus was defined using the Allen Mouse Brain Atlas. In FIJI, a 325 μm x 325 μm square was placed within the LH and a 67826.2μm^2^ ellipse was separately placed within the AH and MPOA to create the regions of interest (ROI) for quantification. The sizes for these ROIs were determined using the scale of the Allen Mouse Brain Atlas and each image had two ROIs per brain region (one ROI/hemisphere). Fiber number (and cell number in the AH) was hand counted in each brain region by a blinded observer. Thresholding was performed on each maximum intensity projection using FIJI for analysis of % area coverage measurement. 1-4 images per animal were quantified for each brain region and measurements for each individual ROI were averaged to provide a single measurement per brain region per animal. OXT-ir and AVP-ir cell bodies found in the AH were quantified following the same procedure as in the PVN.

### qPCR and analysis

#### Tissue collection and punching

At P50-52, animals from a separate cohort of animals that were not exposed to behavioral testing were euthanized with CO_2_. Brains were removed and flash-frozen in 2-methylbutane, then stored at -80°C until sectioning and tissue punching. For tissue punching, brains were mounted in a sterilized cryostat and tissue punches were collected by inserting a sterilized 1.5mm diameter core sampling tool to a depth of 1 mm (Electron Microscopy Sciences). Tissue punches were collected from the NAc, LS, AMY, dHipp and vHipp. These regions were chosen based on their known expression of Oxtr and V1aR, importance to social behavior (Dumais et al., 2013; Smith et al., 2017), and shape limits of the tissue punching procedure. Tissue was collected bilaterally except for the LS. Punches were stored in 500 μL Trizol (Thermo Fisher Scientific) and frozen at -80°C until RNA extraction.

#### RNA extraction

RNA was extracted using phenol-chloroform extraction. Samples were homogenized in 500 μL Trizol (Thermo Fisher Scientific), vortexed at 2000 rpm for 10 min, and rested for 15 min. 100 μL chloroform (Sigma Aldrich) was added and samples were vortexed at 2000 rpm for 2 min and rested for 3 min. Samples were centrifuged at 11,800 rpm for 15 min at 4° C. The aqueous phase was extracted from the phase gradient, 250 µl isopropanol (Sigma Aldrich) and 2 µl Glycogen Blue (Thermo Fisher Scientific) were added to the aqueous phase, and samples were centrifuged at 11,800 rpm for 10 min at 4° C. RNA pellets were rinsed with 500 µl ice-cold 75% ethanol (Thermo Fisher Scientific) twice, centrifuging at 9000 rpm for 5 min at 4° C after ethanol was added each time. Pellets were then re-suspended in nuclease-free (NF) H2O (Fisher Bioreagents) and stored at -80 °C until cDNA synthesis.

#### cDNA synthesis

cDNA synthesis was performed using the QuantiTect Reverse Transcription Kit according to its standard protocol (Qiagen). RNA quantity and quality was assessed using a NanoDrop One C (Thermo Fisher Scientific). Briefly, samples were gDNAse treated in a Thermofisher Applied Biosystems MiniAmp Plus Thermal Cycler for 2 min at 42 °C. Next, Master mix was prepared with 1 µl reverse transcriptase (RT), 4 µl RT buffer, and 1 µl RT primer mix per sample. Master mix was added to suspended RNA. Samples were incubated for 15 min at 42 °C and 3 min at 95 °C. Samples were then diluted to 10 ng/µl and stored at -20°C unless proceeding immediately with qPCR. No-template and no-RT controls were included to verify primer specificity during qPCR.

#### Quantitative PCR (qPCR)

Quantitative PCR (qPCR) was performed using the QuantiNova SYBR Green PCR Kit (Qiagen). PCR primers were designed in-house and purchased from Integrated DNA Technologies (*18S*: F: GAATAATGGAATAGGACCGC, R: CTTTCGCTCTGGTCCGTCTT; *Oxtr*: F: ACCTGGACTCCCACCTATTT, R: GCCGTCTTTCACAAGATACCA; *Avpr1a*: F: ATCCGCACAGTGAAGATGAC, R: GGAATCGGTCCAAACGAAATTG. To prepare the SYBR master mix, 6.5 µl QuantiNova SYBR, 1 µl forward primer, 1 µl reverse primer, and 3.5 µl NF H2O per reaction were combined. 12 µl SYBR master mix per well was plated on MicroAmp Optical 96-well Reaction Plates (Applied Biosystems), then 1 µl template cDNA was added to each well. Samples were run in duplicates. The no-template and no-RT controls generated during cDNA synthesis were run on each plate to verify primer specificity. qPCR was run on a QuantStudio 3 Real-Time PCR machine (Thermo Fisher Scientific). Samples were held at 95 °C for 2 min to activate SYBR. 40 cycles of PCR were performed: samples were held at 95 °C for 5 minutes, ramped to 49.4 or 49.9 °C at a speed of 1.6 °C/sec (for *Oxtr* and *Avpr1a*, respectively), and held at 49.4 or 49.9 °C for 11 sec; then the cycle was repeated. After PCR was complete a melt curve was performed.

### 2-ΔΔCT analysis

Relative gene expression was calculated using the 2-ΔΔCT method, relative to *18*S gene expression and the lowest sample on the plate (Williamson et al., 2011; Livak & Schmittgen, 2001). Microsoft Excel was used for 2-ΔΔCT calculations. Samples were excluded before unblinding if they failed to amplify specifically (i.e. no amplification or multiple melt curve peaks) or if replicates differed by >1 fold change.

### Statistics

All statistical analyses were conducted using GraphPad Prism 10 software. For maternal body weight, a 2-way ANOVA (embryonic day x treatment) was conducted. Litter ratios were analyzed using unpaired t-tests (VEH/CON vs. DEP/MS). Pup body weights, OXT-ir/AVP-ir cells and fibers, and *Oxtr*/*Avpr1a* mRNA were analyzed using 2-way ANOVAs (sex x treatment) followed by Tukey’s posthoc tests. Behavioral outcomes were assessed using mixed effect 2-way ANOVAs (stimulus/chamber [repeated measure] x treatment [between-subjects measure]) followed by Sidak’s multiple comparisons tests. Because males and females were tested using separate apparatuses on different days, male and female behavior results were analyzed separately. Statistical outliers were identified using the ROUT outlier test (Q=1%). Data are expressed as mean ± SEM and statistical significance was set at p < 0.05.

## Results

### Maternal and litter outcomes

While both VEH/CON and DEP/MS dams gained significant weight during pregnancy, this did not differ between treatment groups (see **Table 1** for complete F statistics; **Figure 1B**). DEP/MS exposure had no effect on litter size (**Figure 1C**; t_(12)_=0.28, p=0.79) or on the number of males (**Figure 1D**, t_(12)_=0.56, p=0.59) or females (**Figure 1E**, t_(12)_=0.00, p>0.99) in each litter. Pup body weights were measured at timepoints pre- and post-weaning to assess pup developmental growth trajectory following prenatal DEP/MS. Prior to weaning, at P16, we found an interaction effect; DEP/MS males weighed less than VEH/CON males while DEP/MS females weighed more than VEH/CON females (**Table 1**; **Figure 1F**). At P40, males weighed more than females, consistent with sex differences in size emerging at puberty (**Table 1**; **Figure 1G**). There was also a trend towards a treatment effect with DEP/MS offspring tending to weigh less than VEH/CON offspring (p=0.09).

**Table 1.** 2-way ANOVA results of maternal and offspring body weights. Bolded statistics indicate significant effects (p<0.05) and trends (p<01), P=postnatal day, g = grams.

| Outcome | Treatment | Embryonic Day (dams)<br>or Sex (offspring) | Interaction |
| --- | --- | --- | --- |
| Maternal Body Weight (g) | $F_{(1, 12)} = 0.01, p=0.94$ | **$F_{(1.44, 17.29)} = 460.3, p<0.0001$** | $F_{(5, 60)} = 0.540, p=0.75$ |
| P16 Body Weight (g) | $F_{(1, 86)} = 0.023, p=0.88$ | $F_{(1, 86)} = 0.000, p=0.99$ | **$F_{(1, 86)} = 5.432, p=0.022$** |
| P40 Body Weight (g) | **$F_{(1, 78)} = 2.956, p=0.09$** | **$F_{(1, 78)} = 173.6, p<0.0001$** | $F_{(1, 78)} = 0.688, p=0.41$ |

### Behavior

#### Light-Dark Box Test

In the light-dark box test (**Figure 2A**), we found that DEP/MS did not alter chamber time or fecal boli deposition in males (see **Table 2** for complete F statistics; fecal boli: t _(13)_ = 1.030, p=0.322; **Figure 2B &C**). Males spent more time in the dark side of the box than in the light (*post hoc* light vs. dark: p<0.001) regardless of treatment. In females, there was no main effect of treatment, but a significant treatment x chamber interaction effect (p=0.03) whereby DEP/MS females tended to spend more time in the light (*post hoc* VEH/CON vs. DEP/MS in light chamber: p=0.06) and less time in the dark (*post hoc* VEH/CON vs. DEP/MS in dark chamber: p=0.07) as compared to VEH/CON females. There was also a trend towards an increase in fecal boli deposited in DEP/MS vs. VEH /CON females (fecal boli: t _(15)_ = 1.781, p=0.095; **Figure 2D & E**). VEH/CON females spent more time in the dark side of the box (*post hoc* light vs. dark in VEH/CON females: p<0.001) while DEP/MS females did not (*post hoc* light vs. dark in DEP/MS females: p=0.683).

**Figure 2.**
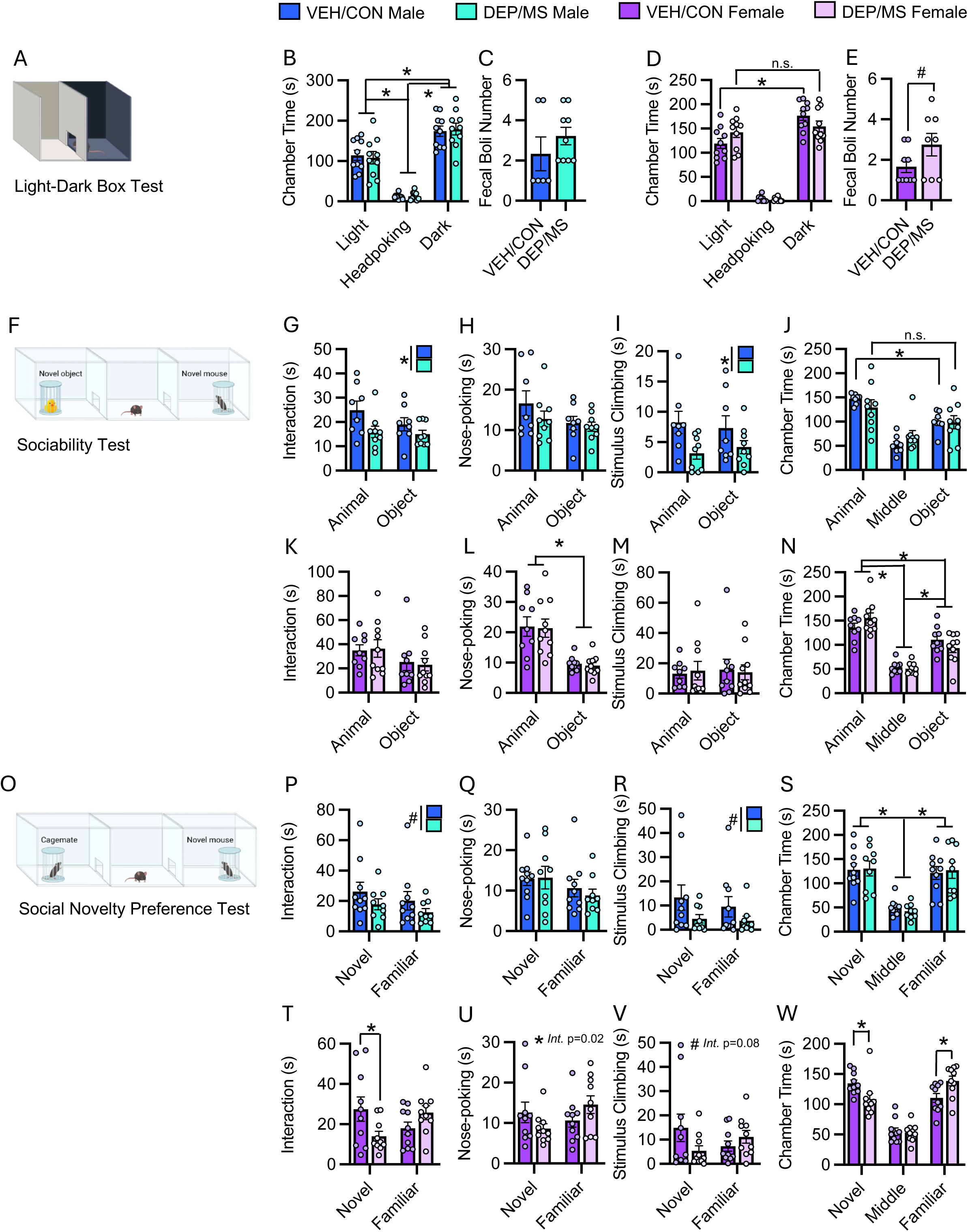
Results of light/dark box and social behavior assays. **A**) Light-Dark Box test. **B** & **D**) Results of light/dark box test in males (**B**) and females (**D**). mixed effects ANOVA (chamber x treatment) with Sidak’s *post hoc* tests. **C**& **E**) Fecal boli deposition in males (**C**) and females (**E**) un-paired t-tests. **F**) Three-chambered sociability assay. **G-I**) In males, DEP/MS reduces interaction interaction time (**G**) and stimulus climbing (**I**) but not nose-poking (**H**). Main effects of treatment in 2-way ANOVAs (stimulus x treatment). **J**) In males, there is a main effect of chamber. *Post hoc* tests reveal that VEH/CON males spend more time in the social chamber than the object chamber, whereas DEP/MS males spend similar times in the social and object chambers. Mixed effects 2-way ANOVA (chamber x treatment) with Sidak’s *post hoc* tests. **K-M**) In females, no effect of DEP/MS or stimulus on interaction (**K**) or stimulus climbing (**M**) but a main effect of stimulus on nose-poking (**L**) 2-way ANOVAs (stimulus x treatment). **N**) In females, there is a main effect of chamber. *Post hoc* tests reveal that females of both treatments spend differing amounts of time in the animal, object, and middle chambers. Mixed effects 2-way ANOVA (chamber x treatment) with Sidak’s *post hoc* tests. **O**) Three-chambered social novelty preference test. **P-R**) In males, DEP/MS tends to reduce interaction (**P**) and stimulus climbing (R) but not nose-poking (**Q**) as compared to VEH/CON. 2-way ANOVAs (stimulus x treatment). **S**) In males, there is a main effect of chamber, with males spending more time in both the novel and familiar chambers than the middle chamber. Mixed effects 2-way ANOVA (chamber x treatment) with Sidak’s *post hoc* tests. T-V) In females, there are interaction effects or trends towards interactions between stimulus and treatment on interaction (**T**), nose-poking (**U**), and stimulus climbing (**V**) with DEP/MS females interacting more with the familiar stimulus and interacting less with the novel stimulus compared to VEH/CON females. 2-way ANOVAs (stimulus x treatment) with Sidak’s *post hoc* tests. **W**) There is an interaction between chamber and treatment on chamber times, with DEP/MS females spending more time in the familiar chamber and spending less time in the novel chamber compared to VEH/CON females. Mixed effects 2-way ANOVA (chamber x treatment) with Sidak’s *post hoc* tests. s = seconds. Data = mean ± SEM, *p<0.05; #p<0.10, Results shown are of main effects unless significant post hoc effects were found in which case those are labeled.

**Table 2.** 2-way ANOVA results of light/dark box, sociability, and social novelty preference tests in males and females. Bolded statistics indicate significant effects (p<0.05) and trends (p<0.1), s: seconds.

| Behavior | Treatment | Stimulus/Chamber | Interaction |
| --- | --- | --- | --- |
| **Light/Dark Box:** |  |  |  |
| <u>Males</u> |  |  |  |
| Chamber time (s) | $F_{(1, 54)} = 0.000, p > 0.99$ | **$F_{(2, 54)} = 105.7, p < 0.0001$** | $F_{(2, 54)} = 0.128, p = 0.88$ |
| <u>Females</u> |  |  |  |
| Chamber time (s) | $F_{(1, 54)} = 0.000, p > 0.99$ | **$F_{(2, 54)} = 197.4, p < 0.0001$** | **$F_{(2, 54)} = 3.504, p = 0.037$** |
| **Sociability:** |  |  |  |
| <u>Males</u> |  |  |  |
| Stimulus interaction (s) | **$F_{(1, 15)} = 5.980, p = 0.02$** | $F_{(1, 15)} = 1.635, p = 0.220$ | $F_{(1, 15)} = 0.884, p = 0.362$ |
| Stimulus investigation (s) | $F_{(1, 15)} = 1.577, p = 0.228$ | $F_{(1, 15)} = 2.524, p = 0.133$ | $F_{(1, 15)} = 0.471, p = 0.503$ |
| Stimulus climbing (s) | **$F_{(1, 15)} = 5.022, p = 0.041$** | $F_{(1, 15)} = 0.002, p = 0.965$ | $F_{(1, 15)} = 0.870, p = 0.366$ |
| Chamber time (s) | $F_{(1, 45)} = 0.001, p = 0.979$ | **$F_{(2, 45)} = 24.370, p < 0.0001$** | $F_{(2, 45)} = 1.423, p = 0.252$ |
| <u>Females</u> |  |  |  |
| Stimulus interaction (s) | $F_{(1, 17)} = 0.009, p = 0.924$ | $F_{(1, 17)} = 2.152, p = 0.161$ | $F_{(1, 17)} = 0.056, p = 0.815$ |
| Stimulus investigation (s) | $F_{(1, 17)} = 0.073, p = 0.790$ | **$F_{(1, 17)} = 25.89, p < 0.0001$** | $F_{(1, 17)} = 0.000, p = 0.985$ |
| Stimulus climbing (s) | $F_{(1, 17)} = 0.002, p = 0.964$ | $F_{(1, 17)} = 0.017, p = 0.899$ | $F_{(1, 17)} = 0.078, p = 0.783$ |
| Chamber time (s) | $F_{(1, 51)} = 0.000, p > 0.999$ | **$F_{(2, 51)} = 65.20, p < 0.0001$** | **$F_{(2, 51)} = 2.506, p = 0.092$** |
| **Social Novelty Preference:** |  |  |  |
| <u>Males</u> |  |  |  |
| Stimulus interaction (s) | **$F_{(1, 17)} = 3.339, p = 0.085$** | $F_{(1, 17)} = 1.078, p = 0.314$ | $F_{(1, 17)} = 0.006, p = 0.939$ |
| Stimulus investigation (s) | $F_{(1, 17)} = 0.207, p = 0.655$ | $F_{(1, 17)} = 1.830, p = 0.194$ | $F_{(1, 17)} = 0.184, p = 0.673$ |
| Stimulus climbing (s) | **$F_{(1, 17)} = 3.661, p = 0.073$** | $F_{(1, 17)} = 0.408, p = 0.532$ | $F_{(1, 17)} = 0.163, p = 0.692$ |
| Chamber time (s) | $F_{(1, 51)} = 0.000, p > 0.999$ | **$F_{(2, 51)} = 28.80, p < 0.0001$** | $F_{(2, 51)} = 0.188, p = 0.830$ |
| <u>Females</u> |  |  |  |
| Stimulus interaction (s) | $F_{(1, 18)} = 0.495, p = 0.491$ | $F_{(1, 18)} = 0.078, p = 0.783$ | **$F_{(1, 18)} = 7.309, p = 0.015$** |
| Stimulus investigation (s) | $F_{(1, 18)} = 0.000, p = 0.990$ | $F_{(1, 18)} = 1.595, p = 0.223$ | **$F_{(1, 18)} = 6.413, p = 0.021$** |
| Stimulus climbing (s) | $F_{(1, 18)} = 0.697, p = 0.415$ | $F_{(1, 18)} = 0.061, p = 0.808$ | **$F_{(1, 18)} = 3.581, p = 0.075$** |
| Chamber time (s) | $F_{(1, 54)} = 0.000, p > 0.999$ | **$F_{(2, 54)} = 63.48, p < 0.0001$** | **$F_{(2, 54)} = 7.283, p = 0.002$** |

#### Sociability Assay

In the sociability assay (**Figure 2F**) in males, there was a main effect of treatment such that total interaction time was lower in DEP/MS as compared to VEH/CON males (main effect of treatment: p=0.027; see **Table 2** for complete statistics; **Figure 2G**). Interaction time is comprised of nose-poking into (investigating) and stimulus climbing on the social or object container. Interestingly, when broken down into its’ component parts, we found that the decrease in interaction time was largely driven by decreased stimulus climbing following DEP/MS (main effect of treatment: p=0.04) rather than nose-poking (main effect of treatment: p=0.23; **Figure 2H & I**). There were no main effects of treatment on chamber time in males. However, there was a main effect of chamber (p<0.001) with more time spent in the social chamber than the middle or novel object chambers (**Figure 2J).** Post hoc comparisons showed that VEH/CON males spent more time in the social chamber as compared to the object chamber (*post hoc* social vs. object in VEH/CON males: p=0.015) while DEP/MS males did not (*post hoc* social vs. object in DEP/MS males: p=0.137). DEP/MS did not affect middle chamber entries in the sociability assay in males (middle chamber entries: t _(15)_ = 0.239, p=0.815).

In females, DEP/MS treatment did not affect stimulus interaction, nose-poking, or stimulus climbing in the sociability assay (**Table 2**; **Figure 2K-M**). There was a main effect of stimulus on nose-poking time, with females investigating the social stimulus more than the object (main effect of stimulus: p<0.001; **Figure 2L**) regardless of treatment. In line with this, there was also a main effect of chamber, with females spending a greater amount of time in the chamber containing the social stimulus than the middle or novel object chambers, regardless of treatment (**Figure 2N**). DEP/MS did not affect middle chamber entries in the sociability assay in females (middle chamber entries: t _(17)_ = 1.191, p=0.250).

#### Social Novelty Preference Test

In the social novelty preference test (**Figure 2O**) in males, DEP/MS tended to decrease interaction time (main effect of treatment, p=0.085; for complete F statistics see **Table 2**; **Figure 2P**) and stimulus climbing time (main effect of treatment, p=0.07; **Figure 2R**). There was no significant effect of DEP/MS on nose-poking (**Figure 2Q**). There was no significant effect of treatment on overall chamber time, but there was a main effect of chamber, with males spending more time in the novel and familiar stimulus chambers than the middle chamber regardless of treatment (**Figure 2S**). DEP/MS tended to reduce middle chamber entries in the social novelty preference test in males, though this did not reach statistical significance (middle chamber entries: t _(17)_ = 1.903, p=0.074).

In females, there was a significant interaction effect on interaction time (stimulus x treatment interaction: p=0.015; **Figure 2T**). Post hoc comparisons revealed a significant decrease in interaction with the novel conspecific following DEP/MS as compared to VEH/CON (*post hoc* VEH/CON vs. DEP/MS novel interaction: p=0.04). This was mirrored in both a significant interaction effect on nose-poking time (stimulus x treatment interaction: p=0.015; **Figure 2U**) and a trend towards an interaction effect on stimulus climbing time (p=0.075; **Figure 2V**) with DEP/MS females shifting towards decreased novel nose-poking/stimulus climbing but more familiar nose-poking/stimulus climbing as compared to VEH/CON females. This shift in preference towards the familiar cage mate was also reflected in chamber times. DEP/MS females spent less time in the novel chamber (*post hoc* VEH/CON vs. DEP/MS novel chamber: p=0.012) and more time in the familiar chamber as compared to VEH/CON females (*post hoc* VEH/CON vs. DEP/MS novel chamber: p<0.01; **Figure 2W**). DEP/MS did not affect middle chamber entries in the social novelty preference test in females (middle chamber entries: t _(18)_ = 0.436, p=0.668).

### OXT and AVP in the Hypothalamus

#### OXT and AVP cell bodies in the PVN and AH

We found that in the more rostral Level 1 PVN sub-region analyzed, DEP/MS tended to increase OXT-ir cell number, regardless of sex as compared to VEH/CON (main effect of treatment p=0.07; see **Table 3** for complete statistics; **Figure 3A&B**). This pattern did not continue more caudally in the Level 2 PVN sub-region, where we did not observe any treatment effects on OXT cell number (**Figure 3C&D**). We observed no significant effects of sex or treatment on AVP-ir cell number in either Level 1 or 2 of the PVN (**Figure 3E-H**), nor did we find any sex or treatment effects on either OXT-ir or AVP-ir cell bodies in the AH which were sparse but present (**Figure 3I-K**).

**Table 3.** 2-way ANOVA results of oxytocin and vasopressin cell counts in males and females. Bolded statistics indicate significant effects (p<0.05) and trends (p<0.1), PVN: paraventricular nucleus of the hypothalamus, AH: anterior hypothalamus.

| Cell Type | Treatment | Sex | Interaction |
| --- | --- | --- | --- |
| **Oxytocin cell bodies:** |  |  |  |
| Level 1 PVN | **$F_{(1, 28)} = 3.547, p=0.070$** | $F_{(1, 28)} = 0.393, p=0.536$ | $F_{(1, 28)} = 0.014, p=0.908$ |
| Level 2 PVN | $F_{(1, 20)} = 0.883, p=0.359$ | $F_{(1, 20)} = 2.438, p=0.134$ | $F_{(1, 20)} = 0.350, p=0.561$ |
| AH | $F_{(1, 28)} = 0.497, p=0.487$ | $F_{(1, 28)} = 0.401, p=0.532$ | $F_{(1, 28)} = 0.026, p=0.873$ |
| **Vasopressin cell bodies:** |  |  |  |
| Level 1 PVN | $F_{(1, 15)} = 0.034, p=0.855$ | $F_{(1, 15)} = 0.014, p=0.901$ | $F_{(1, 15)} = 0.218, p=0.647$ |
| Level 2 PVN | $F_{(1, 19)} = 0.490, p=0.493$ | $F_{(1, 19)} = 0.276, p=0.606$ | $F_{(1, 19)} = 2.267, p=0.149$ |
| AH | $F_{(1, 30)} = 0.000, p=0.984$ | $F_{(1, 30)} = 2.214, p=0.147$ | $F_{(1, 30)} = 0.228, p=0.637$ |

**Figure 3.**
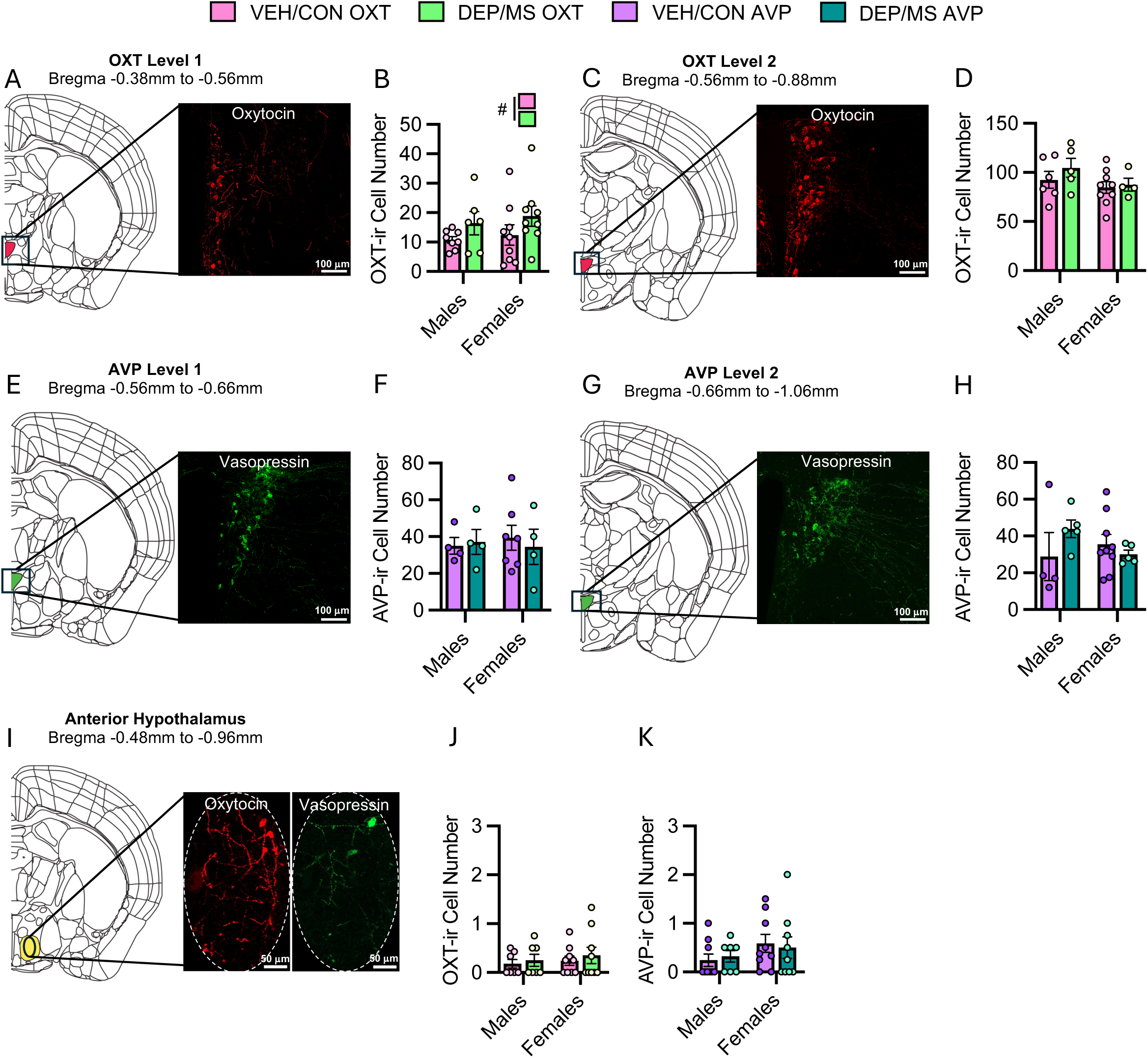
Effects of DEP/MS exposure on OXT-ir and AVP-ir cell number in the PVN. **A**) Representative image of OXT-ir cells in OXT Level 1 in the PVN. **B**) DEP/MS tended to increase OXT-ir cell number in both males and females in OXT Level 1. **C**) Representative image of OXT-ir cells in OXT Level 2. **D**) No effect of DEP/MS or sex on OXT-ir cell number in OXT Level 2. **E**) Representative image of AVP-ir cells in AVP Level 1. **F**) No effect of DEP/MS or sex on AVP-ir cell number in AVP Level 1. **G**) Representative image of AVP-ir cells in AVP Level 2 **H**) No effect of DEP/MS or sex on AVP-ir cell number in AVP Level 2. **I**) Representative images of OXT-ir and AVP-ir cells in the AH. **J**) No effect of DEP/MS or sex on OXT-ir cell number in the AH. **K**) No effect of DEP/MS or sex on AVP-ir cell number in the AH. 2-way ANOVAs (sex x treatment). Data = mean ± SEM, #p<0.1.

#### OXT and AVP fibers in the LH, AH, and MPOA

We also assessed OXT-ir and AVP-ir fiber density in three hypothalamic nuclei that play varying roles in social behavior: the LH, AH, and MPOA. We used two measures to assess fiber density: fiber number and % area covered by fibers. We did not observe any significant effects of treatment or sex in any of the regions analyzed on either of these measures (for complete F statistics see **Table 4**; **Figure 4A-O**). Interestingly, we did find a trend towards a sex difference in OXT-ir fiber number in the LH with females having more fibers than males (main effect of sex: p=0.09) but this did not reach statistical significance.

**Figure 4.**
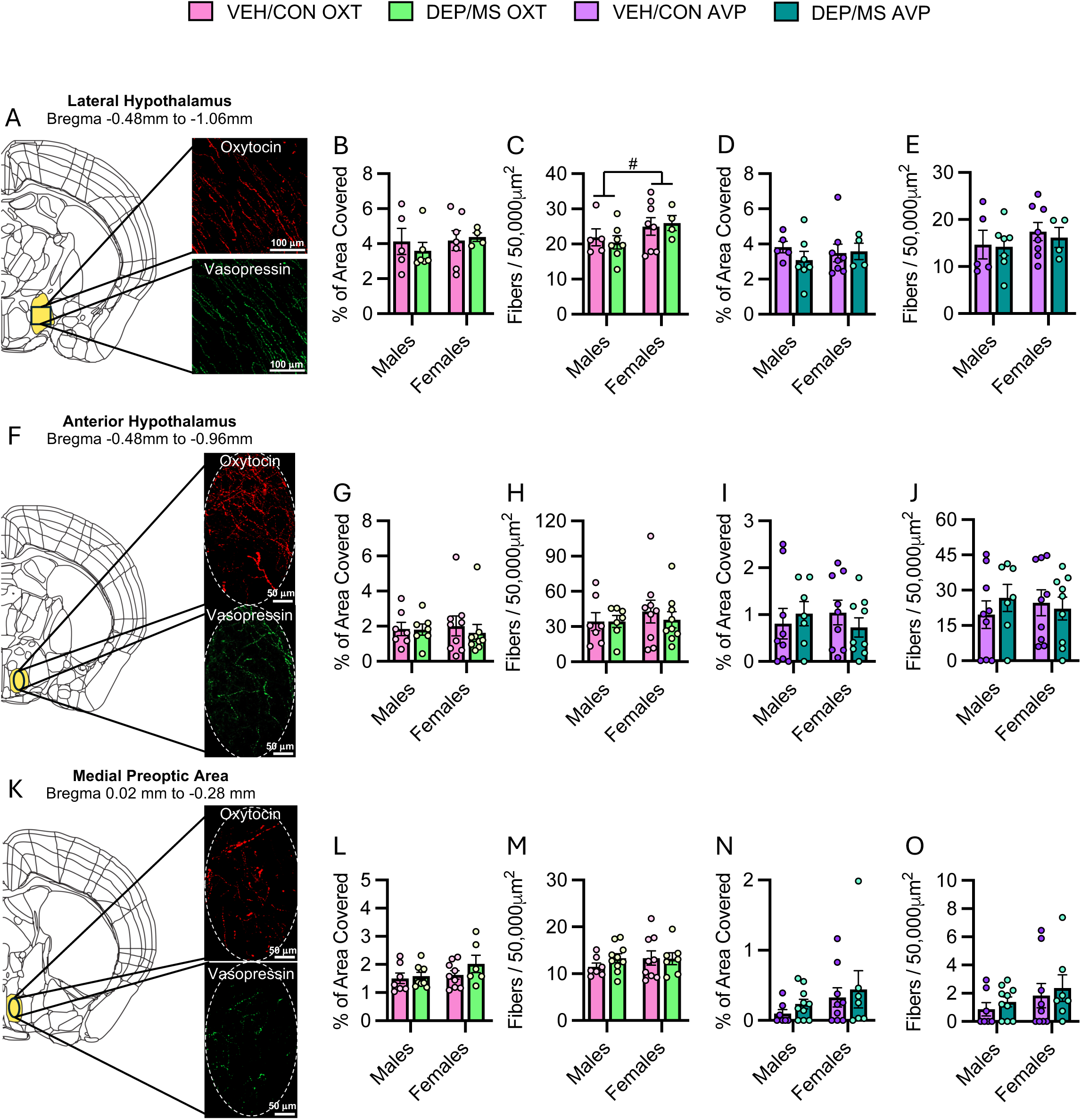
Effects of DEP/MS exposure on OXT-ir and AVP-ir fibers in the hypothalamus. **A**) Representative images of OXT-ir and AVP-ir fibers in the LH. **B-E**) No effect of DEP/MS or sex on OXT-ir or AVP-ir fibers in the LH, but a trend towards a sex difference in OXT-ir fiber number (**C**). **F**) Representative images of OXT-ir and AVP-ir fibers in the AH. G-J) No effect of DEP/MS or sex on OXT-ir or AVP-ir fibers in the AH. **K**) Representative images of OXT-ir and AVP-ir fibers in the MPOA. **L-O**) No effect of DEP/MS or sex on OXT-ir or AVP-ir fibers in the MPOA. 2-way ANOVAs (sex x treatment). Data = mean ± SEM, #p<0.1.

**Table 4.**
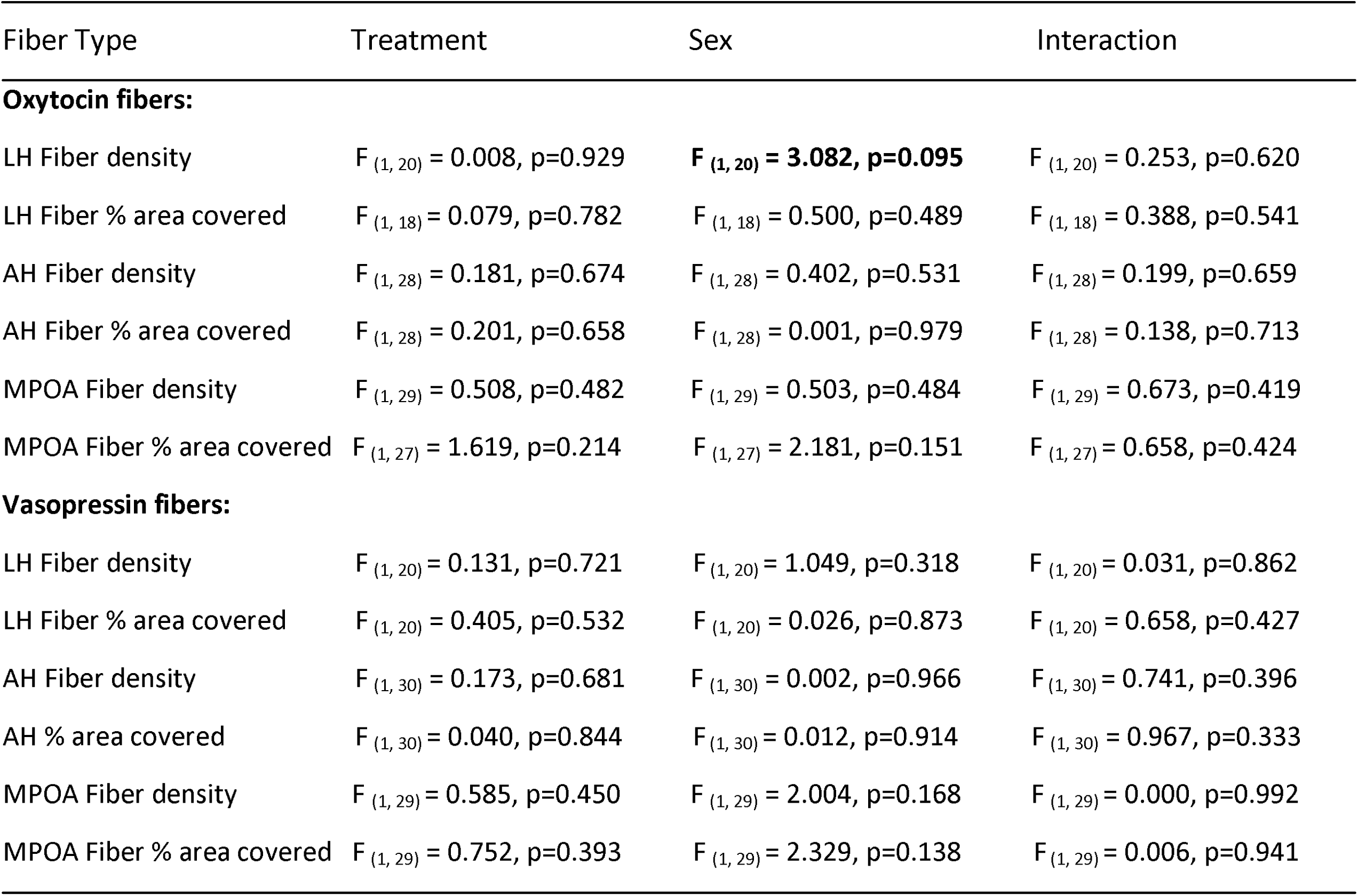
2-way ANOVA results of oxytocin and vasopressin fiber counts in males and females. Bolded statistics indicate significant effects (p<0.05) or trends (p<0.1), LH: lateral hypothalamus, AH: anterior hypothalamus, MPOA: medial preoptic area of the hypothalamus.

### *Oxtr* and *Avpr1a* gene expression in regions of the SDMN

We observed numerous effects of sex and treatment on *Oxtr* and *Avpr1a* mRNA expression (for complete F statistics see **Table 5**). In the NAc, males exhibited greater *Oxtr* mRNA expression compared to females (main effect of sex: p<0.01), but there was no effect of DEP/MS on *Oxtr* mRNA expression (Figure **5A**). Interestingly, DEP/MS increased *Avpr1a* mRNA expression in both males and females in the NAc as compared to VEH/CON (main effect of treatment: p=0.015, **Figure 5A**). In the LS, females expressed more *Oxtr* and *Avpr1a* mRNA than males (main effects of sex: *Oxtr* p=0.048, *Avpr1a* p=0.009), but there was no effect of DEP/MS (**Figure 5B**). In the AMY, there were no significant effects of sex or treatment on *Oxtr* mRNA expression (**Figure 5C**). However, there was a trend towards a decrease in *Avpr1a* mRNA following DEP/MS as compared to VEH/CON (main effect of treatment p=0.086, **Figure 5C**). Additionally, females exhibited greater *Avpr1a* mRNA expression than males (main effect of sex: p=0.036, **Figure 5C**). In the dHipp, neither treatment nor sex altered *Oxtr* mRNA expression (**Figure 5D**). There was a trend towards a significant increase in *Avpr1a* mRNA expression following DEP/MS as compared to control (main effect of treatment: p=0.05), as well as a trend towards greater *Avpr1a* mRNA expression in females compared to males (main effect of sex p=0.06, **Figure 5D**). Finally, in the vHipp, there were no effects of treatment or sex on *Oxtr* or *Avpr1a* mRNA expression (**Figure 5E**).

**Figure 5.**
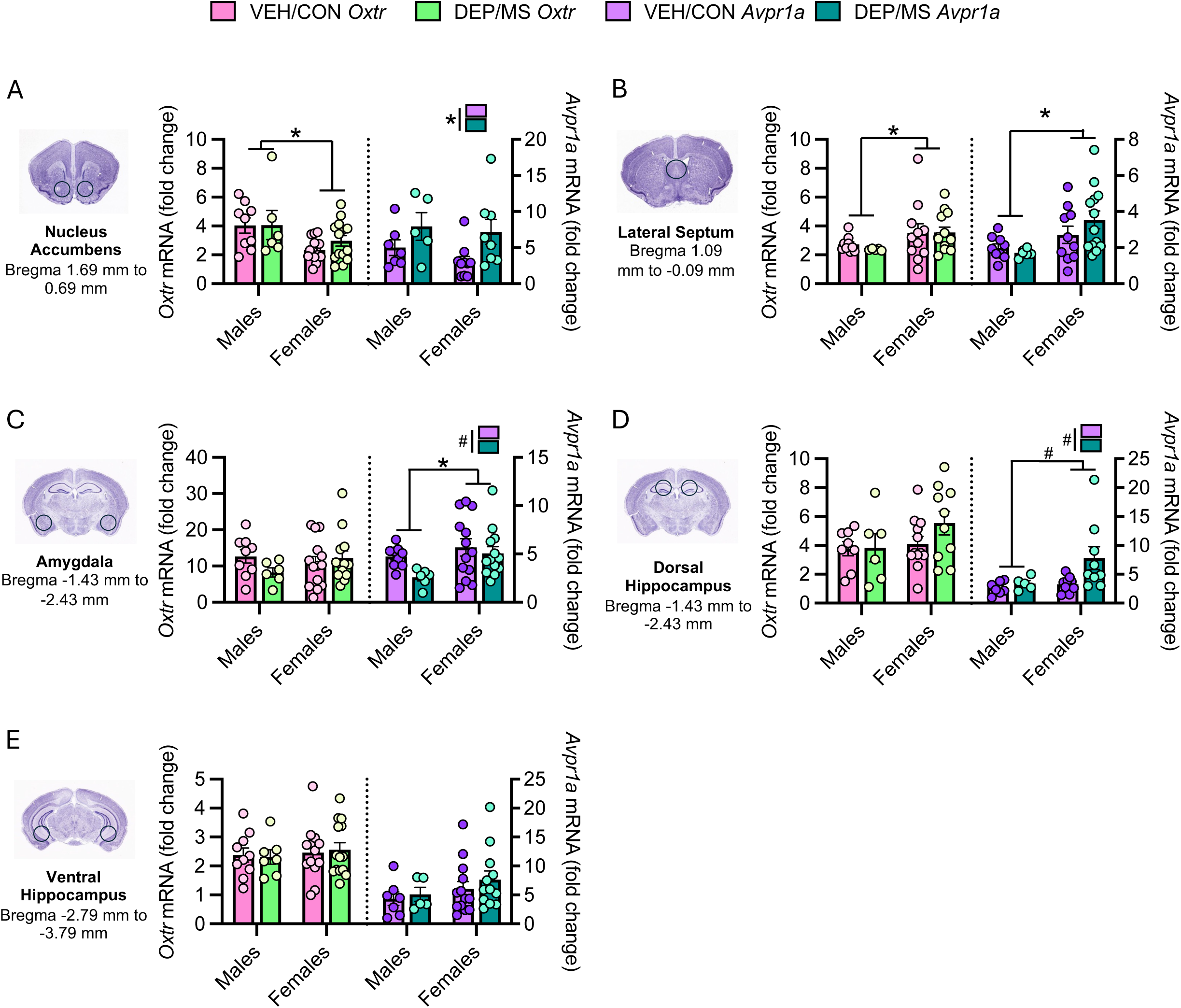
Effects of DEP/MS exposure on OXT-ir and AVP-ir fibers in the hypothalamus. **A**) In the NAc, males express more *Oxtr* mRNA than females. DEP/MS increases *Avpr1a* mRNA expression in the NAc in both males and females. B) No effect of DEP/MS on *Oxtr* or *Avpr1a* mRNA expression in the LS, though females express more *Oxtr* and *Avpr1a* mRNA than males. **C**) In the AMY, there is no effect of DEP/MS or sex on *Oxtr* mRNA expression. Females express more *Avpr1a* mRNA than males in the AMY, and there is a trend towards decreased *Avpr1a* mRNA expression following DEP/MS. **D**) In the dHipp, there is no effect of DEP/MS or sex on *Oxtr* mRNA expression. DEP/MS tends to increase *Avpr1a* mRNA expression in both males and females and females tend to express more *Avpr1a* mRNA than males. **E**) No effect of DEP/MS or sex on *Oxtr* or *Avpr1a* mRNA expression in the vHIPP. 2-way ANOVAs (sex x treatment). Data = mean ± SEM, *p,0.05, #p<0.1.

**Table 5.** 2-way ANOVA results of Oxtr and Avpr1a expression in males and females. Bolded statistics indicate significant effects (p<0.05) and trends (p<0.1), NAc: nucleus accumbens, LS: lateral septum, AMY: amygdala, dHIPP: dorsal hippocampus, vHIPP: ventral hippocampus.

| Brain Region | Treatment | Sex | Interaction |
| --- | --- | --- | --- |
| ***Oxtr* expression:** |  |  |  |
| NAc | $F_{(1, 40)} = 0.491, p=0.488$ | **$F_{(1, 40)} = 8.993, p=0.005$** | $F_{(1, 40)} = 0.436, p=0.513$ |
| LS | $F_{(1, 34)} = 0.103, p=0.750$ | **$F_{(1, 34)} = 4.229, p=0.048$** | $F_{(1, 34)} = 1.168, p=0.684$ |
| AMY | $F_{(1, 38)} = 0.572, p=0.454$ | $F_{(1, 38)} = 0.285, p=0.600$ | $F_{(1, 38)} = 2.060, p=0.159$ |
| dHIPP | $F_{(1, 32)} = 1.143, p=0.293$ | $F_{(1, 32)} = 2.056, p=0.161$ | $F_{(1, 32)} = 0.911, p=0.347$ |
| vHIPP | $F_{(1, 40)} = 0.003, p=0.954$ | $F_{(1, 40)} = 0.379, p=0.542$ | $F_{(1, 40)} = 0.087, p=0.770$ |
| ***Avpr1a* expression:** |  |  |  |
| NAc | **$F_{(1, 26)} = 6.787, p=0.015$** | $F_{(1, 26)} = 0.968, p=0.334$ | $F_{(1, 26)} = 0.192, p=0.665$ |
| LS | $F_{(1, 34)} = 0.343, p=0.562$ | **$F_{(1, 34)} = 7.739, p=0.009$** | $F_{(1, 34)} = 1.491, p=0.231$ |
| AMY | **$F_{(1, 37)} = 3.120, p=0.086$** | **$F_{(1, 37)} = 4.755, p=0.036$** | $F_{(1, 37)} = 0.960, p=0.334$ |
| dHIPP | **$F_{(1, 26)} = 4.134, p=0.052$** | **$F_{(1, 26)} = 3.798, p=0.062$** | $F_{(1, 26)} = 2.046, p=0.165$ |
| vHIPP | $F_{(1, 33)} = 0.558, p=0.460$ | $F_{(1, 33)} = 1.914, p=0.176$ | $F_{(1, 33)} = 0.073, p=0.790$ |

## Discussion

Taken together, our results indicate several important findings. First, in accordance with previous work, DEP/MS exposure does not alter maternal weight gain during pregnancy or overall litter composition but does appear to subtly impact offspring weight gain during the early postnatal period. Second, in contrast to our hypothesis of male-specific phenotypes following DEP/MS, we observed effects of DEP/MS exposure on social behavior in both male and female offspring, with different specific social outcomes impacted in each sex. Finally, while DEP/MS exposure does not appear to significantly impact OXT-ir or AVP-ir cell or fiber density in the hypothalamus, our results show that DEP/MS exposure alters *Avpr1a* mRNA but not *Oxtr* mRNA expression in several brain regions: it increases *Avpr1a* mRNA expression in the NAc and tends to increase expression in the dHipp, while tending to decrease expression in the AMY, in both sexes.

### Divergent effects of DEP/MS exposure on social behavior in males and females

We assessed the impact of DEP/MS exposure on several behavioral assays (light/dark box, sociability, and social novelty preference) during the adolescent period in both male and female offspring. Interestingly, we observed effects of DEP/MS exposure on behavior in both sexes, although the specific behavioral outcomes impacted differed between males and females. DEP/MS exposed males spent less time engaged in overall interaction and stimulus climbing as compared to their VEH/CON counterparts in the sociability assay and tended to spend less time in engaging in these behaviors in the social novelty preference test. The decrease in interaction time in the sociability assay in males following DEP/MS was more pronounced for the social stimulus as compared to the object, but this did not reach statistical significance. DEP/MS exposed females, on the other hand, exhibited a shift in their social preference towards the familiar cage mate in the social novelty preference test, as evidenced by reduced interaction, nose-poking, and chamber times associated with the novel stimulus as compared to VEH/CON females. Our findings in the sociability assay in males are only somewhat in line with previous work (Block et al., 2022; Smith et al., 2023). We did observe a decrease in social interaction in male offspring following DEP/MS, however object interaction was also decreased. Moreover, we found that this decrease in total interaction was largely driven by a decrease in climbing of the stimuli rather than nose-poking into them. Interestingly, while VEH/CON males spent significantly more time in the social chamber of the test, DEP/MS males did not. Together, these findings may reflect a more general decrease in exploratory behavior than previously reported. The decrease in climbing behavior observed following DEP/MS exposure could reflect changes in either exploratory motivation or complex motor abilities (also disrupted in neurodevelopmental disorders; Monteiro et al., 2022; Ji et al., 2023) and this should be the subject of future investigations.

Contrary to our hypothesis that we would see male-specific social deficits, here, we also find decreases in social novelty preference in female offspring following DEP/MS exposure. Interestingly, effects on social behavior in females in this model have been observed previously, but at an earlier developmental timepoint. Block et al., (2022) measured ultrasonic vocalizations, a measure of social communication at P8 and found effects of DEP/MS exposure in both sexes on number of calls and total time spent calling, which supports the idea that female social development is also impacted by DEP/MS exposure. Decreases in social behavior in females have also been observed in other models of prenatal air pollution or stress exposure in rodents. For example, studies of prenatal through postnatal exposure to air pollutants have found decreases in sociability, social novelty preference, and altered ultrasonic vocalizations in both males and females (Zhou et al., 2021; Chang et al., 2018). Prenatal maternal restraint stress also impairs social behavior in both male and female offspring in adulthood (Gur et al., 2019; Chen et al., 2024). These findings are important given that neurodevelopmental disorders are male-biased but not male-specific; females are also diagnosed. Further, current animal models of behavior and neurodevelopmental disorders were mostly developed using male subjects only and are likely not adequately capturing female-specific mechanisms and behaviors (Shansky, 2024; Becegato & Silva, 2024; Smith & Kingsbury et al., 2020).

In our previous work, we found that social behavior deficits induced by DEP/MS exposure in males were accompanied by changes in the composition of the gut microbiome (Smith et al., 2023). Furthermore, these social behavior deficits were prevented by cross-fostering at birth, a manipulation that restored the gut microbiome to the control phenotype. The rodent gut microbiome is known to shift depending on factors including housing conditions, institution, diet, and vendor (Franklin & Ericsson, 2017; Long et al., 2021). While we designed our experiments so that bedding, diet, and other conditions were consistent with previous studies, these experiments were conducted at a new institution. It would be interesting to determine whether the changes we observe here in behavior in both sexes are accompanied by changes in the gut microbiome and whether this is a factor that might at least partially explain the differing behavioral results. The gut-brain axis has been implicated in the regulation of social behavior in numerous studies (Desbonnet et al., 2014; Hsiao et al., 2013; Holder et al., 2019; Buffington et al., 2016; Sherwin et al., 2019; Sgritta et al., 2019; Lu et al., 2018; Arnold et al., 2025) and there is emerging evidence that sex differences exist in the composition of gut microbiota, in both humans and mice (Kim et al., 2020). C57Bl/6J mice—the mouse strain used in the present study—exhibit some of the most robust sex differences in gut microbiota, including expression of the genus *Bacteroides*, with greater abundance in females than males (Org et al., 2016). *Bacteroides* is differentially expressed in individuals with ASD (Sharon et al., 2019) and decreased in abundance in males exposed to DEP/MS (Smith et al., 2023). One possibility is that differences in the abundance of bacterial genera such as *Bacteroides* between sexes, exposures (such as DEP/MS), and studies may contribute to sex differences in social behavior outcomes.

In the light-dark box test, we observed no main effect of treatment, but DEP/MS exposed females spent slightly more time in the light side of the box compared to VEH/CON females and tended to deposit more fecal boli during behavior, which is often interpreted as a measure of anxiety (Kulesskaya et al., 2014; Contet et al., 2001; Rodgers et al., 2002). This is the first study to assess behavior in the light-dark box test following prenatal DEP/MS exposure. However, previous findings in the DEP/MS model have been mixed with respect to the assessment of anxiety-like behavior using other assays. Smith et al., (2023) found no differences in behavior in the open field test following DEP/MS exposure during adolescence (Smith et al., 2023). In contrast, in adulthood, Bolton et al. (2013) found that DEP/MS increased anxiety-like behavior in the elevated zero maze in both male and female offspring (Bolton et al., 2013). More sensitive and/or naturalistic assays of anxiety-like behavior, such as the looming-shadow task (Bolton et al., 2022), may be helpful in clarifying this behavioral dimension.

### No significant effects of DEP/MS exposure on OXT-ir or AVP-ir cell/fiber density in the hypothalamus

Following behavioral testing, offspring brains were collected to determine whether there are long-lasting developmental changes to OXT/AVP innervation following DEP/MS that corresponded to behavioral effects. We found that DEP/MS exposure had no significant effects on OXT-ir or AVP-ir cell number in the PVN or AH, nor on fiber density in the AH, LH, or MPOA. We observed a trend towards higher OXT-ir cell number in the more anterior portion of the PVN assessed, but this did not reach statistical significance (p=0.07). Still, this finding is interesting given that OXT-ir cell number has been shown to increase in the PVN with manipulations to the gut microbiome such as treatment with *L. reuteri* (Varian et al., 2023). The present study is the first to quantify OXT and AVP innervation following prenatal air pollution or DEP/MS exposure, and these systems have been much more thoroughly characterized following perinatal exposure to other environmental toxicants, the effects of which on OXT and AVP vary considerably depending on dosage, timing of exposure, sex and other factors (for review see Martin et al., 2025). For example, Reilly et al., (2022) also found no overall effects of prenatal exposure to the endocrine disruptor Aroclor 1221 (a polychlorinated biphenyl) on either OXT or AVP cell number in the PVN, but the suggestion of a rostro-caudal shift (Reilly et al., 2022). DEP/MS exposure induces an MIA cascade in dams (Block et al., 2022) and our findings are largely in line with those of two studies of OXT and AVP expression in the hypothalamus following MIA, both in rats (Taylor et al., 2012; Breach et al., 2024). Taylor et al., (2012) used in situ hybridization to assess *AVP* mRNA in the PVN, supraoptic nucleus (SON) and superchiasmatic nucleus (SCN) of the hypothalamus and found no effects of maternal lipopolysaccharide (LPS)-induced immune activation on *AVP* mRNA expression in any of these nuclei, although AH, LH, and MPOA were not assessed (Taylor et al. 2012). Similarly, Breach et al., (2024) used an allergic asthma-induced MIA model and found no effects of MIA on OXT-ir or AVP-ir cell or fiber density in the PVN, SON, but found a male-specific decrease in AVP-ir fiber density in the LH; MPOA and AH were not assessed (Breach et al., 2024). Interestingly, both studies did observe effects, particularly on AVP, in extrahypothalamic regions such as the medial amygdala (MeA) and bed nucleus of the stria terminals (BNST; Taylor et al., 2012; Breach et al., 2024). These effects will be discussed further in the context of our receptor expression findings. A limitation of this investigation is that while we have assessed *Oxtr* and *Avpr1a* mRNA in extrahypothalamic regions we did not include analysis of OXT-ir or AVP-ir cell or fiber densities in regions such as the LS, AMY, or BNST. Therefore, it is possible that DEP/MS impacts OXT and AVP expression in these regions and should be the subject of future investigations.

### DEP/MS effects on *Avpr1a*, but not *Oxtr*, mRNA in regions of the SDMN

We observed treatment effects on *Avpr1a* mRNA expression and sex effects on both *Avpr1a* and *Oxtr* mRNA expression. Most notably, *Avpr1a* mRNA expression was significantly increased in the NAc in both males and females following DEP/MS. This is especially interesting given that the NAc is a brain region in which numerous changes have been observed following DEP/MS exposure (Smith et al., 2023), including male-specific decreases in dopamine receptor expression and shifts in microglial morphology and gene expression. This finding is also in line with Breach et al., (2024) which observed higher AVP-ir fiber density in the NAc after MIA, albeit only in male offspring. There is limited work indicating the functional relevance of changes to V1aR expression in the NAc in mice. Vasopressin acting at V1aRs in the NAc and ventral pallidum has been shown to mediate social behaviors such as partner preference formation in monogamous prairie voles (Hammock & Young, 2006). For example, the formation of a partner preference induces V1aR upregulation in the NAc of female prairie voles and partner preference formation is prevented by V1aR antagonist administration (Wang et al., 2013). This is relevant given that females exposed to DEP/MS display an increased preference for the familiar cage mate along with higher NAc-*Avpr1a* expression. One possibility is that higher NAc-*Avpr1a* following DEP/MS exposure serves to increase familiar rather than novel interactions. *Oxtr* mRNA was not impacted by DEP/MS exposure, but we did observe a sex difference in the NAc with higher *Oxtr* mRNA in males as compared to females. This sex difference is consistent with previous autoradiography work in rodents showing higher Oxtr binding in males than in females (Dumais et al., 2013).

*Avpr1a* mRNA also tended to increase in the dHipp and decrease in the AMY following DEP/MS exposure as compared to VEH/CON. Very little previous work has characterized the impacts of prenatal air pollution and/or stress on hippocampal V1aR expression in either sex. In typically developed rodents, vasopressin signaling at hippocampal V1b receptors is critical for social recognition and social aggression (Leroy et al., 2018; Cilz et al., 2019). One study of maternal restraint stress during pregnancy found that *Avpr1a* mRNA in the hippocampus of female offspring was positively correlated with amount of maternal care they received (Schmidt et al., 2018). Still, much remains unknown as to what the implications of higher *Avpr1a* mRNA in females following DEP/MS are for behavioral outcomes. In contrast to this finding, we observed a trend towards decreased *Avpr1a* mRNA expression in the AMY. While air pollution/stress effects on AMY V1aR have not previously been characterized, perinatal exposure to the pesticide chlorpyrifos and the plasticizer bisphenol A have both been shown to alter AVP and V1aR in the MeA (Venerosi et al., 2015; Goldsby et al., 2017; Arambula et al., 2018). Similarly, Breach et al. (2024) found higher AVP-ir fiber density in the MeA in both males and females following MIA. In contrast, Taylor et al., (2012) found lower *AVP* mRNA expression in the AMY following a different, LPS-induced MIA. V1aRs in the AMY regulate social behaviors including social recognition in the MeA (Nephew & Bridges, 2008; Arakawa et al., 2010) and maternal aggression in the central amygdala (Caughey et al., 2011; Oliveira et al., 2022). One limitation of our approach is that our tissue punches of the AMY include all subregions (medial, central, and basolateral). Therefore, specific investigation of receptor mRNA and/or protein within the various subregions of the AMY and extended amygdala is an important avenue for future research.

We found no effects of treatment on *Oxtr* or *Avpr1a* mRNA in the LS or vHipp. However, we found sex differences in both *Oxtr* or *Avpr1a* mRNA in the LS with higher expression in females than in males for both. These sex differences reflect previously observed sex differences in these receptors in the LS. Specifically, Smith et al., (2017) found higher Oxtr binding density in the dorsal LS and higher V1aR binding density in the intermediate LS in female rats as compared to male rats using receptor autoradiography. This in line with the previously discussed sex difference in Oxtr in the NAc, which is also consistent with previous work using receptor autoradiography (Dumais et al., 2013). A limitation of the present study is the quantification of mRNA which does not necessarily translate to the protein level. However, the consistency of our findings with previous work using receptor autoradiography supports the idea that our observed effects on mRNA could be indicative of protein level changes. Future work will aim to characterize differences in V1aR in the NAc and AMY using methods that directly quantify protein expression, such as with the use of reporter mouse lines.

### Conclusions

In conclusion, our results have several important implications. First, our finding of behavioral changes following prenatal exposure to DEP/MS in both males and females underscores the need to strongly consider the effects of neurodevelopmental insults on females in addition to males. Our observation of different behavioral effects in each sex following DEP/MS highlights the need to expand our behavioral repertoire to fully capture male- and female-specific responses to prenatal toxicant and stress exposures. We also find specific effects of prenatal DEP/MS exposure on *Avpr1a* mRNA expression in multiple brain regions, in the absence of robust changes in *Oxtr* mRNA or changes in hypothalamic OXT-ir and AVP-ir cells or fibers. Overall, this work suggests that prenatal exposures to combined air pollutants and maternal stress have long-term impacts on behavior and V1aR expression in the social brain.

## Acknowledgments

The authors would like to thank Bret Judson for support with microscopy, Ehri Ryu for support with statistical analysis, and the animal care staff at Boston College for the laboratory animal maintenance.

## Funding

This work was supported by the National Institute of Environmental Health Sciences (NIEHS) grant R00ES033278 to CJS.

## Author contributions

MCS and CJS designed the study and wrote the paper. MCS, EMM, and CJS analyzed the data. MCS performed the DEP/MS exposures and collected the maternal and litter outcome data. MCS, EMM, CJS, JX, and JTB performed the behavioral experiments and MCS, JTB, JX, and SB conducted behavioral scoring. MCS performed immunohistochemistry and MCS, JTB, and JX conducted OXT-ir and AVP-ir cell and fiber quantification. EMM, MJK, NYL, and MFW extracted RNA, made cDNA, and performed qPCR and analysis. CJS supervised all phases of the project.

## Notes

### Competing Interest Statement

The authors have declared no competing interest.

